# Reliability of canopy photography for forest ecology and biodiversity studies

**DOI:** 10.1101/2024.08.29.610276

**Authors:** Anouk von Meijenfeldt, Francesco Chianucci, Francesca Rigo, Jente Ottenburghs, Andreas Hilpold, Marco Mina

**Affiliations:** Institute for Alpine Environment, Eurac Research, Drususallee/Viale Druso 1, I-39100 Bolzano, Italy; Forest Ecology and Forest Management Group, Wageningen University & Research, Bronland 10, 6708 Wageningen, The Netherlands; CREA, Research Centre for Forestry and Wood, Viale Santa Margherita 80, 52100 Arezzo, Italy; Department of Agricultural, Forest and Food Sciences (DISAFA), University of Turin, 10095 Grugliasco, Italy; Wildlife Ecology and Conservation Group, Wageningen University & Research, Bronland 10, 6708 Wageningen, The Netherlands

**Keywords:** canopy photography, understory vegetation, forest structure, leaf area index, hemispherical photography, mountain forests

## Abstract

Understory is a key component of forest biodiversity. The structure of the forest stand and the horizontal composition of the canopy play a major role on the light regime of the understory, which in turn affects the abundance and the diversity of the understory plant community. Reliable assessments of canopy structural attributes are essential for forest research and biodiversity monitoring programs, as well as to study the relationship between canopy and understory plant communities. Canopy photography is a widely used method but it is still not clear which photographic techniques is better suited to capture canopy attributes at stand-level that can be relevant in forest biodiversity studies. For this purpose, we collected canopy structure and understory plant diversity data on 51 forest sites in the north-eastern Italian Alps, encompassing a diversity of forest types. Canopy images were acquired using both digital cover (DCP) and hemispherical (DHP) photography. Canopy structural attributes were then compared to tree species composition data to evaluate whether they were appropriate to differentiate between forest types. Additionally, we tested what canopy attributes derived from DCP and DHP best explained the species composition of vascular plants growing in the understory. We found that hemispherical canopy photography was most suitable to capture differences in forest types, which was best expressed by variables such as leaf inclination angle and canopy openness. On our sites, DHP-based canopy attributes were also able to better distinguish between different conifer forests. Leaf clumping was the most important attribute for determining plant species distribution of the understory, indicating that diverse gap structures create different microclimate conditions enhancing diverse plant species with different ecological strategies. This study supports the reliability of canopy photography in forest ecology and biodiversity monitoring, but also provide insights for increasing understory diversity in managed forests of high conservation value.

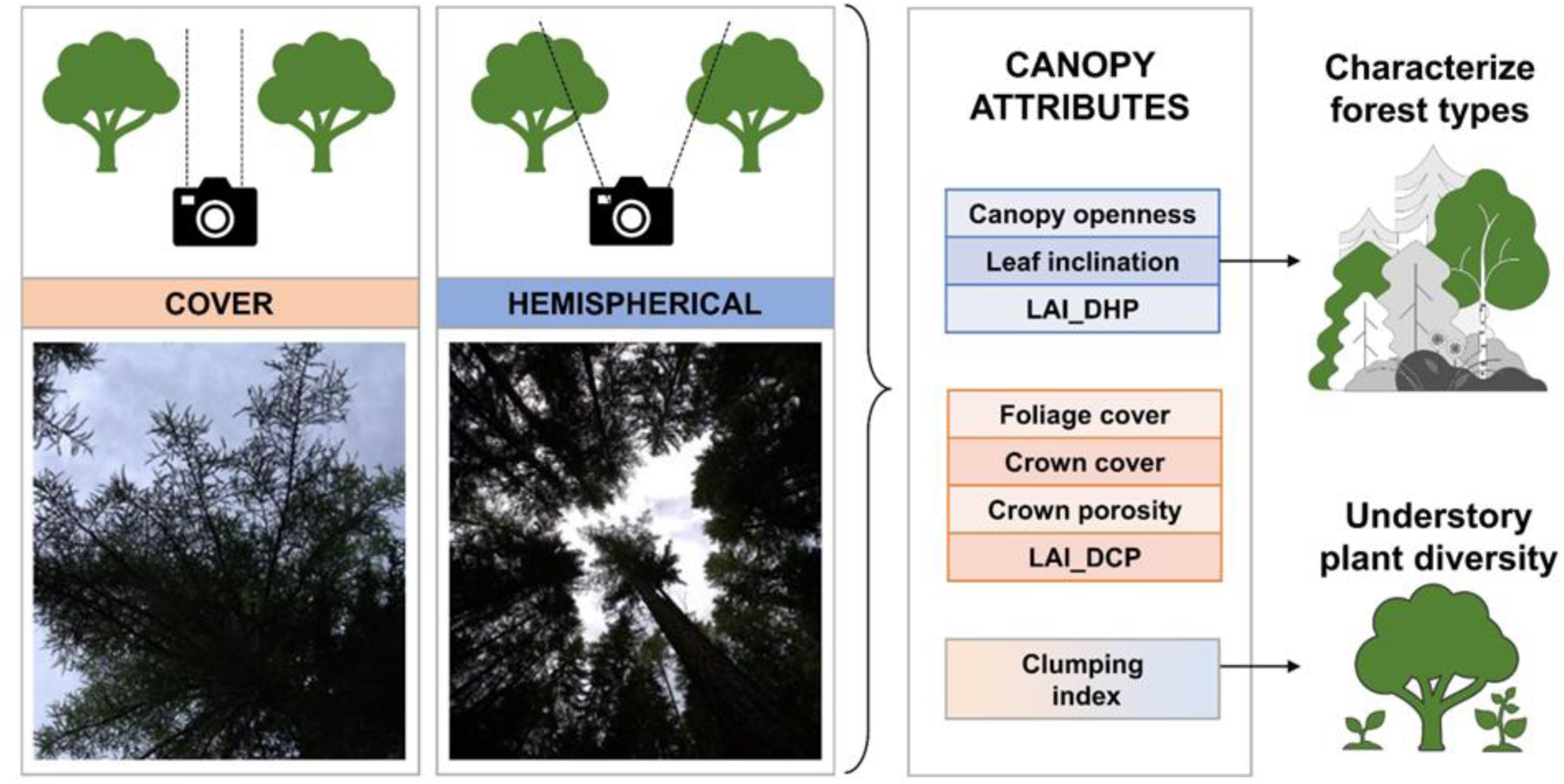

## 1. Introduction

The structure of the forest canopy plays a crucial role in sustaining biodiversity (Nakamura et al., 2017). A well-developed canopy creates a complex vertical structure, offering various habitat and niches for forest dwelling species, from insects and birds to mammals and epiphytes (Rigo et al., 2024). Canopy heterogeneity enhances species richness by providing diverse microhabitats and resources, which reduce competition and promote coexistence (Ishii et al., 2004). Canopy structure also has a direct effect on the light regime of the understory, which in turn affects the abundance and the diversity of the understory plant community (Tinya et al., 2009), which is a crucial component of forest biodiversity (Grime, 1998). Thus, precise assessments of canopy structural attributes are essential in both forest research and ongoing biodiversity monitoring programs (Nakamura et al., 2022). These measurements provide key data for better understanding and managing forest ecosystems effectively.

Among different methods, canopy photography is a widely used one to characterize canopy structural attributes (Chianucci, 2020; Li et al., 2023). This method involves capturing the canopy structure by pointing a photo camera towards the zenith below the trees and deriving the proportions of sky, leaves, and gaps between the leaves from the picture. From the estimated gap fraction, canopy attributes like leaf area index, canopy openness, and crown cover are calculated by applying theoretical gap fraction formulas (Chianucci, 2020). Although canopy photography is an established method (Hill, 1924), it has gained increasing popularity in the last decades due to advances in digital photography technology, which yielded decreased costs of camera equipment and easier processing of pictures with open-source tools (Chianucci et al., 2022; Chianucci and Macek, 2023). Recently, canopy photography has also been used to link canopy structure to forest biodiversity, including understory vegetation (Depauw et al., 2020) and the functional diversity of bats and birds (Rigo et al., 2024). Digital hemispherical photography (DHP) is amongst the most widely used canopy photographic technique (Fournier and Hall, 2017). DHP has the advantage of capturing the largest footprint of the canopy in a single image using fisheye lenses, allowing to infer canopy light regime and leaf area index (Hederová et al., 2023). However, a main drawback of the method is the perceived sensitivity of results to image acquisition and processing (for a review, see Chianucci 2020). An alternative, a more recent technique called digital cover photography (DCP) was introduced by Macfarlane et al (2007) and consists of acquired restricted images with a digital camera and a normal (35 mm equivalent) lens, which yields an approximately 30° field of view. This method has recently gained popularity since it achieves higher-resolution images of the canopy and is relatively insensitive to image acquisition (camera exposure), while image processing is simpler than DHP (Macfarlane et al., 2007). Restricted photography is mostly used for capturing canopy cover, but it also provides additional canopy structure attributes such as foliage clumping and crown porosity. Its main limitation compared to DHP is that it requires assumption on leaf angle distribution to derive leaf area index (LAI) from canopy cover (Chianucci, 2020; Macfarlane et al., 2007). As both methods have advantages and disadvantages, understanding which approach is most suitable for answering specific forest ecology questions would support their operational use.

Canopy attributes are important for ecologically characterising forest vegetation (e.g., conifer *vs* broadleaved *vs* mixed or late seral *vs* light-demanding dominated forests), as different tree species have different crown characteristics, which lead to variation in forest canopy structure (Thomas et al., 2016). These include different leaf angle orientation, which is a rather species-specific attribute (Pisek et al., 2022). In addition, the value of LAI is related to climate and plant functional type at geographically-relevant scale (Parker, 2020). Temperate shade tolerant forest tree species like beech (*Fagus sylvatica*) or holm oak (*Quercus ilex*) may display higher LAI values, with almost non-transparent crowns, which can allow to discriminate these forests from e.g. boreal forests, which typically are dominated by larger, between-crowns gaps (Nilson, 1999). However, very few attempts have been made to ecologically-characterize forest vegetation based on canopy structure. Depauw et al. (2020) identified research plots based on similarity of soil nutrient availability, water holding capacity, and land-use history, while Hederová et al. (2023) selected their research plots based on the similarity of tree species composition. Using directly measured canopy structural attributes for ecologically-based selection of forest sampling plots could be more efficient than relying on other subjective classification of forest composition, while also ensuring comparability between research outcomes and facilitating the distinction between the forest plots. It is not clear which canopy structure parameters are optimal for characterizing forest types, neither what photographic method (e.g., DHP vs DCP) might be more appropriate for forest sampling.

Another important component of the forest that is influenced by light conditions and canopy structure is the community of understory plants (Ádám et al., 2018; Hederová et al., 2023; Sercu et al., 2017). Several studies have shown that canopy structure impacts the distribution and diversity of understory growth (Ádám et al., 2018; Ewald, 2000; Hederová et al., 2023; Lu et al., 2013; Ou et al., 2015; Tinya et al., 2009; Yu and Sun, 2013). However, there has not been a consensus on the extent of canopy impact nor the significance of light structure on plant diversity. For instance, Lu et al. (2013) reported a weak correlation between herbaceous species richness and light flux, while Ou et al. (2015) found that the combination of factors, including soil organic matter, soil pH, overstory Shannon diversity, and total solar radiation transmitted by the canopy could explain 99% of the variation of the understory diversity. Ou et al (2015), however, also found that light parameters, apart from total solar radiation transmitted by the canopy, such as canopy openness, were only weakly correlated with the understory diversity. This contradicts the studies of Yu and Sun (2013) and Hederová et al. (2023) which demonstrated that canopy openness is an important predictor of species in the understory. Although several of these studies demonstrated an effect of canopy structure on understory plant communities, it is still important to determine which structural elements are most influential, and consequently, which canopy photography method might be more appropriate to provide the most relevant parameters.

In this study, we evaluated whether canopy attributes can be used to discriminate different forest vegetation types, testing whether two different techniques – hemispherical (DHP) and cover canopy (DCP) photography – can be used for forest biodiversity studies. By working on multiple forest types across a mountain region in the Italian Alps, we aimed at answering the following research questions:

1. What canopy photography method and canopy attributes are most suitable to capture differences in forest types?
2. What canopy attributes best explain the species composition of understory plants?

For the first question, we evaluated whether canopy attributes derived from two different canopy photography methods can be used to differentiate forest types across a large environmental gradient. For the second question, we explored the predictive power of canopy properties on richness and composition of understory vascular plants. We hypothesized that (H1) hemispherical photography is more suitable for distinguishing forest types as this method is better suited to capture canopy attributes at stand-level (Fournier and Hall, 2017; Hederová et al., 2023). We also hypothesized that (H2) canopy openness, being the variable more directly related to light availability at the forest floor (Beeles et al., 2021; Berdugo and Dovciak, 2019), is the most important attribute for explaining differences in understory plant variation.

## 2. Materials and methods

### 2.1 Study area

The study was carried out within the Autonomous Province of Bolzano-South Tyrol (hereafter South Tyrol), a 7400 km^2^ mountainous province in northern Italy. South Tyrol is located at the northernmost point in Italy and entirely situated in the Alps. The provincés landscape has a wide elevational range, spanning from 204 m a.s.l. at the southern valley bottom to 3905 m a.s.l. at its highest peak. Forests cover almost half of the provincés surface, which due to its complex topography accommodates a large variety of forest types, from low-elevation manna ash-hop hornbeam (*Fraxinus ornus*, *Ostrya carpinifolia*) mixed with oaks (*Quercus pubescens* and *Quercus petraea*) to European beech stands (*Fagus sylvatica*) often mixed with silver fir (*Abies alba*) and Norway spruce (*Picea abies*). The latter is the most common species forming pure and productive forests covering the montane and the subalpine belt, where spruce communities transition towards European larch (*Larix decidua*) and Swiss stone pine (*Pinus cembra*) until the upper timberline.

### 2.2 Forest and understory plant data

We collected data on forest structure and understory plants on 51 forest sampling plots within the entire study area (Fig. 1a). Sampling plots were selected used a random stratification approach in the framework of the Biodiversity Monitoring South Tyrol programme (Hilpold et al., 2023; Rigo et al., 2024) in order to cover the main forest types of the region (Autonomous Province of Bolzano/Bozen, 2010).

**Figure 1.**
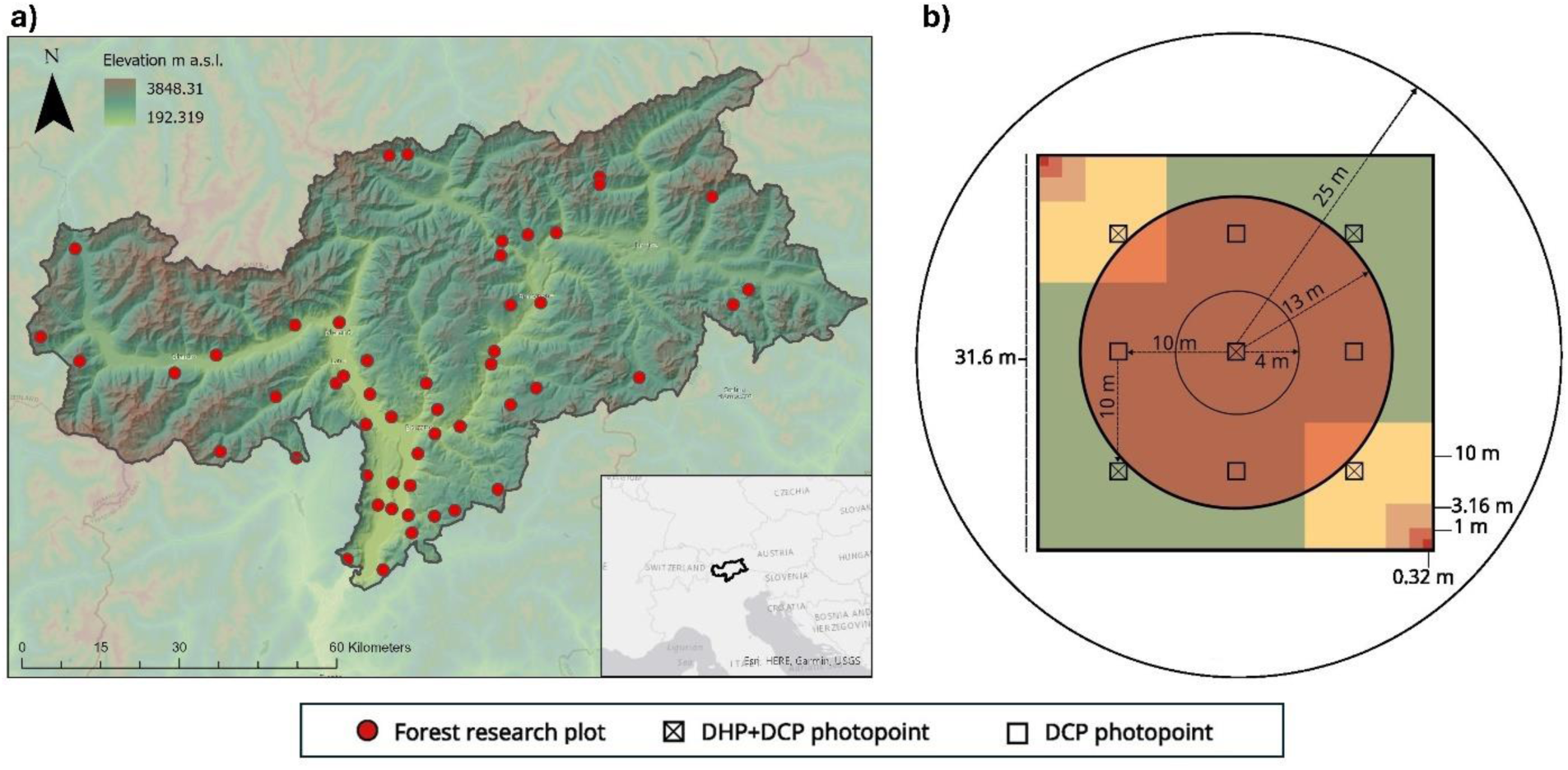
Left: forest sites across the entire study area of South Tyrol. Right: scheme of the photo points within the sampling plot. The understory plant survey plot (green square) with the nested subplots (yellow, orange red squares) is portrayed underneath the forest inventory plot (concentric circles 25-m, 13-m, 4-m) used for collecting forest structure and composition.

Forest structure and compositional data was collected on all sites between 2021 and 2023. Site, management and overstory descriptive characteristics were collected on a ground reference area of 2000 m^2^ (25-m radius from the centre of the sampling plot), while standing trees were measured on two concentric plots (13-m and 4-m radius from the centre, Fig.1b) following the sampling protocol of the Italian National Forest Inventory (Gasparini et al., 2022). None of the sampling plots had signs of recent (<5 years) harvesting interventions. To make sure that no silvicultural intervention took place in the research plots in the last 10 years, we also retrieved and checked the forest management plans of the associated forest stand. Species abundance of the understory plants was collected on the same sites between 2019 and 2023 following the botanical survey protocol of the Biodiversity Monitoring South Tyrol (Hilpold et al., 2023). This vegetation survey was executed on a monitoring plot of 1000 m^2^ (31×31 m), with two subplots of 100 m^2^ (10×10 m) on the upper left and lower right corners. These 100 m2 plots were then divided in a nested design, in which vascular plants were determined with increased level of accuracy. In a series of four nested plots, each square is 10 percent of area from the next square. In the smallest plot the species were counted to estimate abundance per species, in the three larger plots the cover percentage of other species not present in the smaller plots were estimated. For a detailed description of the botanical sampling protocol see Hilpold et al. (2023). A species abundance matrix was made with this data by taking the mean percentages of the two subplots and extrapolating these percentages to the whole 1000 m^2^ monitoring plot. Since we targeted our analysis on herbaceous plant species, we excluded shrub species and tree saplings from the final dataset of understory plants, with exception of small shrub species under 1 m of height, such as blueberries (*Vaccinium myrtillus*), raspberries and brambles (*Rubus spp.*), purple broom (*Chamaecytisus purpureus*) and butchers’ broom (*Ruscus aculeatus*).

### 2.3 Canopy photography data

Digital canopy photographs were collected in a systematic grid within each plot (Figure 1b) using a Sony A6000 camera. Two different methods were used to collect canopy images: cover (DCP) and hemispherical (DHP) photography. DCP images (Macfarlane et al., 2007) were acquired using a 50 mm narrow lens (SELP-1650, E PZ 16-50 mm, f3.5-5.6) resulting in a restricted 30° field of view centred at the zenith (Figure 2). DHP images were obtained with the camera equipped with a full-frame fisheye lens (Walimex Pro 8 mm f/2.8 UMC Fisheye II E), capturing the whole zenith angular range across the diagonal (Figure 2). Due to the different focal length, nine pictures were collected for DCP, and five pictures were collected for DHP (Figure 1b). Images were acquired early in the morning, or in late afternoon, in diffuse light conditions, minimizing the effect of direct sunlight in the images. This approach also allowed for the best contrast between sky and canopy, facilitating subsequent image classification steps (Chianucci, 2020). Aperture was set to F-8 and applying a relative exposure value on two stops of underexposure (REV -2), checking the exposure with the camera histogram. The pictures were shot in RAW mode. RAW images were pre-processed using the *bRaw* package (Chianucci, 2022) to convert it into single channel (blue only) jpeg images and applying a linear contrast stretch to pixel values. This procedure allows reducing the influence of photographic exposure on canopy images (for details, see Macfarlane et al. (2014) and (Chianucci, 2022)).

**Figure 2.**
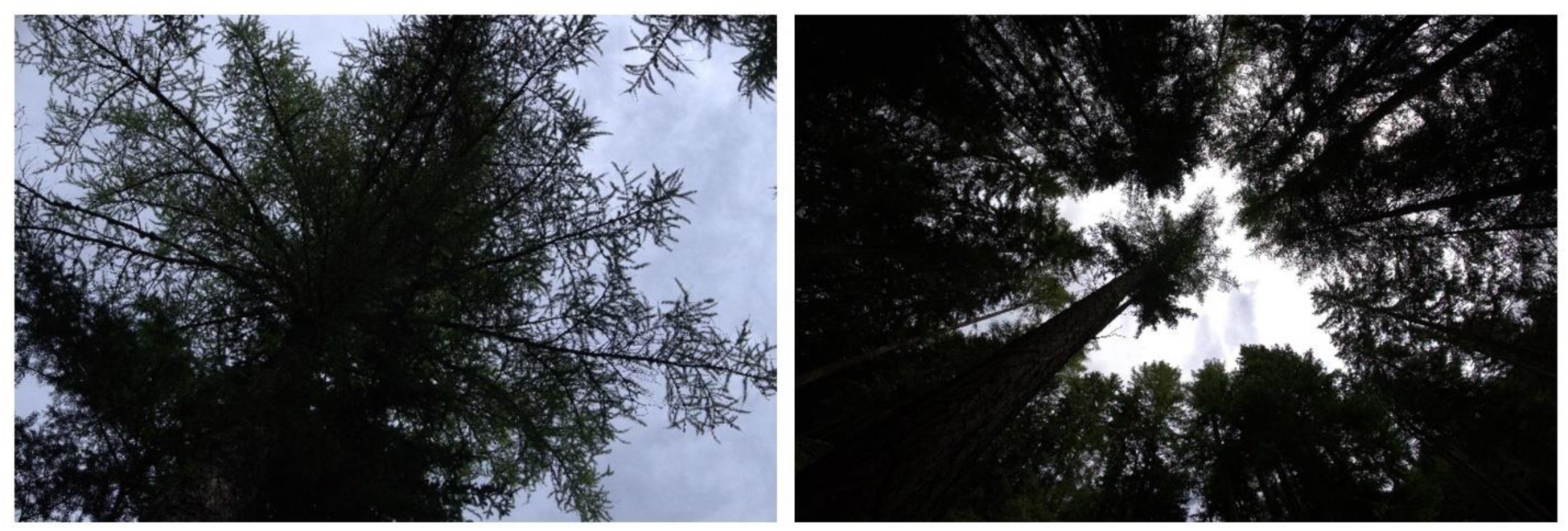
Two canopy photos taken at the same photo point, on the left with restricted view lens (DCP) and on the right with the hemispherical lens (DHP).

The structural attributes of the canopy (Table 1) were then inferred from DCP and DHP by processing canopy images using respectively the *coveR* (Chianucci et al., 2022) and *hemispheR* (Chianucci and Macek, 2023) packages in R. In both cases, the analysis involved thresholding the single, blue channel, to get a binary image of canopy (0) and sky pixels (1), from which the gap fraction can be calculated. For DCP, gaps are further separated into large, between-crown gaps and small, within-crown gaps, to calculate two canopy cover attributes (foliage and crown; see Figure A1), and correct leaf area indices for clumping (Figure A2). For DHP, the package *hemispheR* also allows correcting for lens distortion. Gap fraction was calculated for 7 zenith rings, each 10° in size, and 8 azimuth segments. Effective and actual LAI were calculated using the Miller (1967) theorem, while two clumping indices were calculated from respectively a finite-length averaging method (LX; Lang and Xiang, 1986)) and an ordered weighted log average method (LXG; Chianucci et al., 2019)). Canopy openness (CO) and mean leaf inclination angle (LI, in degrees, using the Elliposidal method by Campbell, 1986) were also calculated.

**Table 1.** Canopy structural attributes collected with either hemispherical lens (DHP) or a restricted view lens (DCP) and their meaning.

| DHP |  | DCP |  |
| --- | --- | --- | --- |
| **LAIe_dhp** | Effective LAI | **FC** | Foliage Cover |
| **LAI_dhp** | Actual LAI | **CC** | Crown Cover |
| **CI_dhp** | Clumping Index | **CP** | Crown Porosity |
| **CI1_dhp** | Alternative Clumping Index 1 | **LAIe_dcp** | Effective LAI |
| **CI2_dhp** | Alternative Clumping Index 2 | **LAI_dcp** | Actual LAI |
| **CO** | Canopy Openness | **CI_dcp** | Clumping index |
| **LI** | Mean Leaf Inclination |  |  |

### 2.4 Statistical tests and analysis

We performed multiple statistical tests to assess the correlation of the DHP and DCP canopy structural attributes. We used k-means clustering with the algorithm of Hartigan and Wong (1979) from the *stats* package in R to categorize forest types based on their canopy structure. To visualize the clustering, we performed a Principal Component Analysis (PCA), with forest research plots as points and canopy structures as vectors. After performing the cluster analysis, we checked the actual tree species composition – derived from inventory sampling in the field – to define whether the clustering based on canopy structure was effective in separating forest types, and which DCP or DHP canopy attributes were more suitable to capture such differences. We applied Kruskal-Wallis tests to check for significant differences between DHP and DCP cluster groups in terms of canopy structural attributes but also in terms of species richness and Shannon diversity of understory plants.

We also conducted a canonical correlation analysis (CCA), using the canopy structural attributes as the environmental elements, the understory plant species abundance as explanatory variable, and the forest plots as dependent variable. A CCA is a restricted correlation analysis and explains which explanatory variables have the highest relative importance in explaining the species composition of understory vascular plants. We selected this method because the species data showed a gradient of 7.9 standard deviations, so a linear method was not appropriate. We used the Bray-Curtis method to calculate the distance between plots because it accounts for zero-inflated species abundance data (Bray and Curtis, 1957). We chose the relevant structural attributes based on backward selection of parameters which were significantly explaining species variation, and highly collinear parameters (based on variance inflation factors, VIF) were removed from the model. The remaining parameters (clumping index from DCP and DHP, and the leaf inclination) had a VIF lower than 5 and significantly explained species variance. Random Monte Carlo permutations (499) were run to verify whether the resulting model was significantly better than a random model. We used Canoco 5 (Jiangshan, 2013), a software package aiding multivariate analysis of ecological data, to extract the sum of constrained eigenvalues and divide it by the sum of all eigenvalues to obtain the percentage explained per canopy structure. All analyses were conducted in the R language and environment for statistical computing v4.4.1 (R Core Team, 2024). Specifically, we used the packages *vegan* (Oksanen et al., 2024) for computing species diversity indices and *tidyverse* (Wickham et al., 2019) for data processing and visualisation.

## 3. Results

Among the 30 tree species surveyed within the 51 research plots, Norway spruce (*Picea abies*) was the most common, making up 21.0% of the total amount of recorded trees (Table A1). Oak species (*Quercus petraea, pubescens*) followed with 16.8%, then hop hornbeam (*Ostrya carpinifolia*) at 13.2%, European larch (*Larix decidua*) at 11.4%, Scots pine (*Pinus sylvestris*) at 6.8%, and European beech (*Fagus sylvatica*) at 5.6%.

### 3.1 Characterizing forest types with different canopy photography methods

The cluster analysis performed on the canopy structural attributes grouped the forest sites in four classes for DHP and three classes for DCP (Figure 3a-b). The optimal number of clusters was determined by analysing different measures of cluster validation, among them the within-cluster sum of squares and the silhouette index (Rousseeuw, 1987). The optimal number of clusters were chosen based on where the decrease of total within sum of squares seemed to stagnate, and where the silhouette index was highest (Figure A3). The canopy attributes from DHP were more effective in grouping forest sites according to their actual species composition (Figure 3c) than those derived from DCP, for which the separation according to tree species composition was less evident (Figure 3d). For the DHP method, the two most important canopy structure parameters for discriminating forest types were leaf inclination angle (LI) and canopy openness (CO; Figure 3a). For the DCP method, the most important canopy structure parameter was leaf area index, either with (LAI) or without (LAIe) correction for clumping (Figure 3b). For both methods, the first PCA axis explained almost all variation (DCP: 98.5%, DHP: 85.8%). Generally, these results showed that canopy attributes derived from DHP performed better than those derived from DCP to characterize differences among forest types.

**Figure 3.**
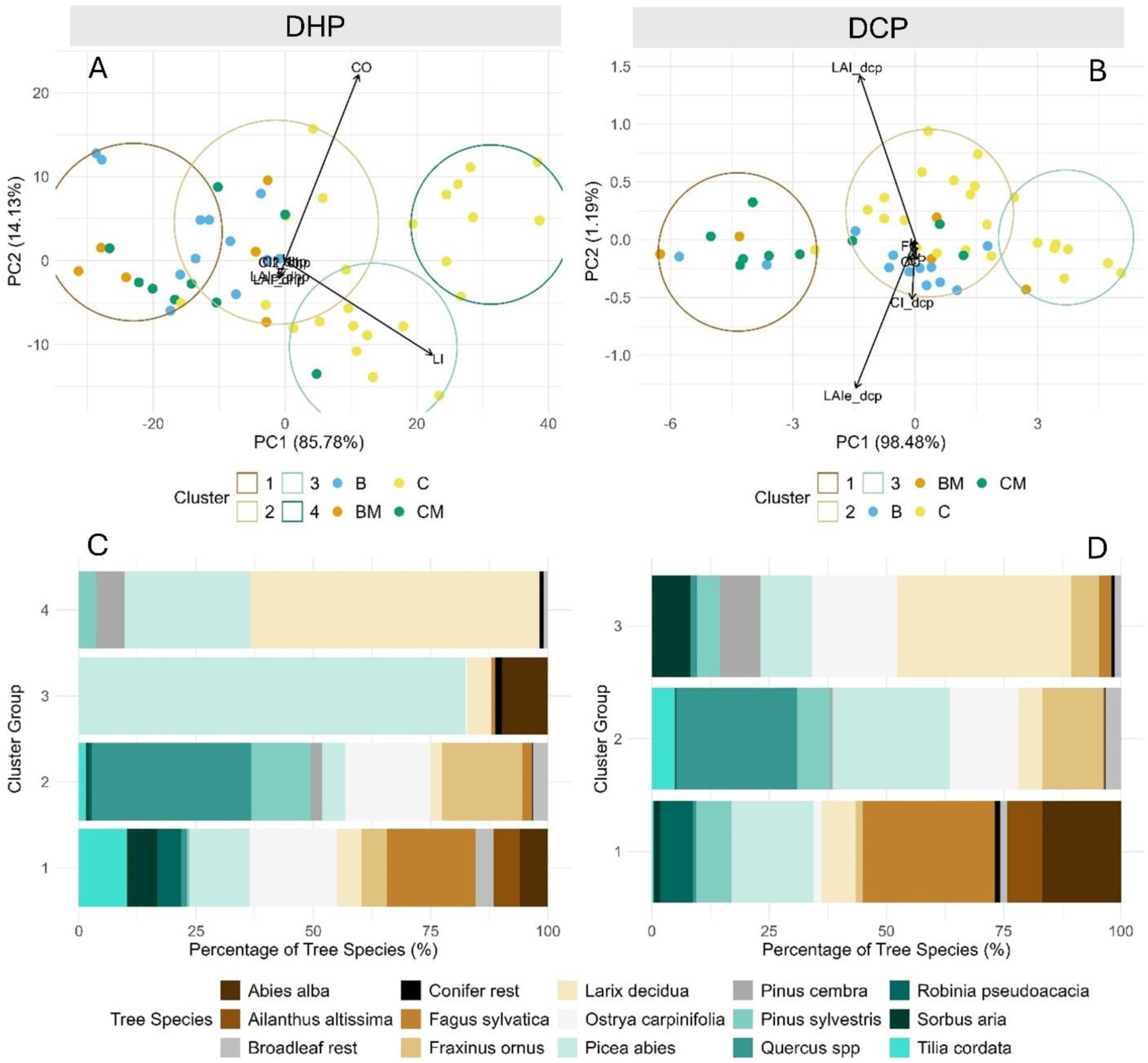
Upper panels (a, b): PCA’s with forest sampling plots as points and DHP/DCP canopy structural elements as vectors. The forest sampling plots have been circled according to their k-means clusters, while the colour of the points indicate the percentage of mixing between broadleaves and conifers (B = broadleaves plot with less than 10% of conifeŕs basal area; BM = broadleaves/mixed plot with basal area of conifers between 10 and 50%; CM = conifer/mixed plot with conifeŕs basal area between 50 and 90%; C = conifer plot with >90% basal area conifer). Lower panels (c, d): percentage of tree species, as derived from field sampling, within the forest plots clustered according to DHP (panel c) and DCP (panel d) canopy variables.

The four groups clustered with DHP attributes differed significantly according to levels of canopy openness (Kruskal-Wallis test, χ2 = 34.8, p-value < 0.001) and leaf inclination (Kruskal-Wallis test, χ2 = 43.6, p-value < 0.001; Figure 4). Despite showing a less evident separation according to species composition, the three groups clustered with DCP attributes also differed significantly according to actual leaf area index (Kruskal-Wallis test, χ2 = 39.0, p-value < 0.001) and effective leaf area index (Kruskal-Wallis test, χ2 = 39.0, p-value < 0.001; Figure A4).

**Figure 4.**
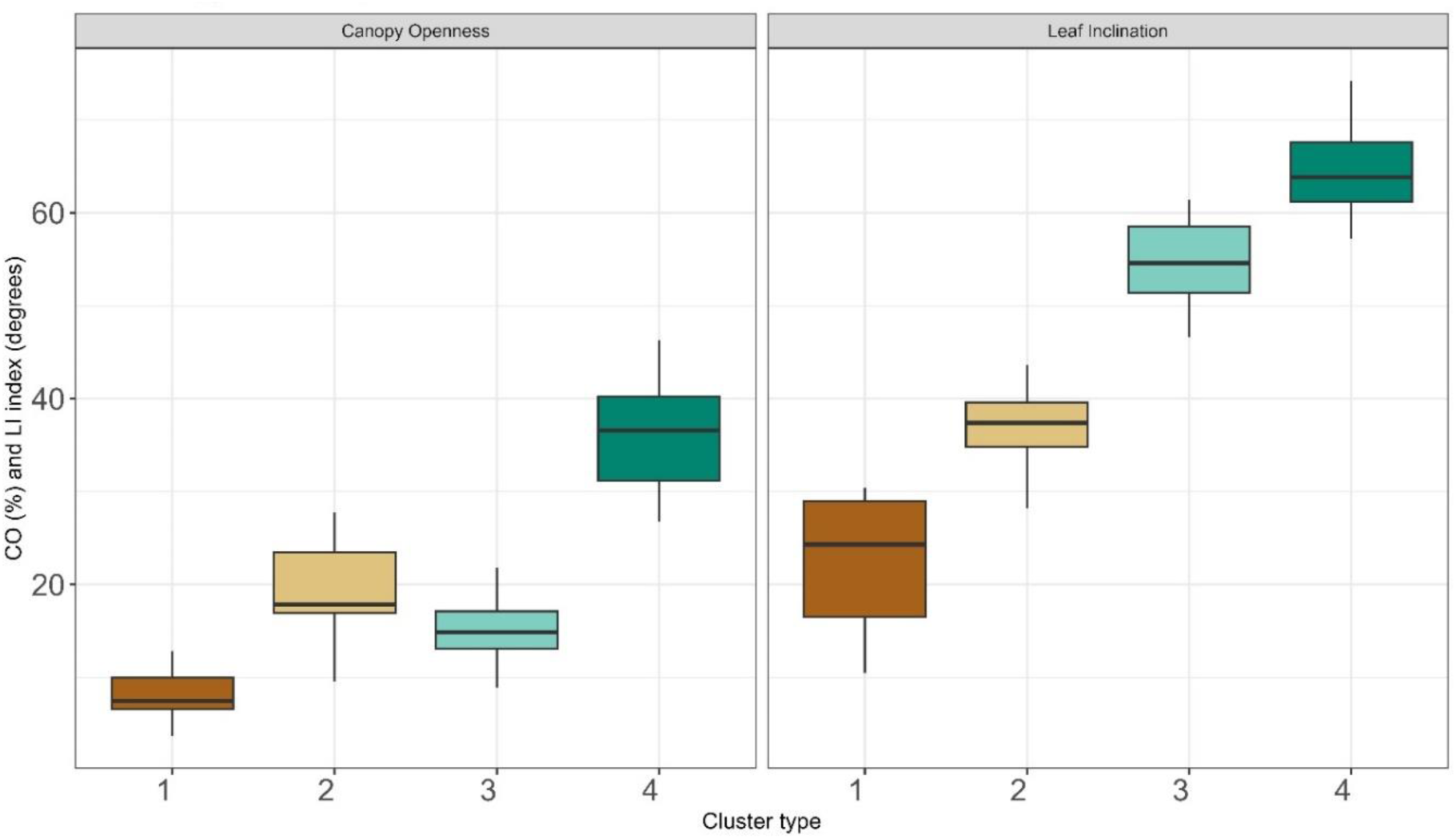
Differences between the four DHP cluster groups for the two canopy attributes canopy openness (left) and mean leaf inclination angle (right). As statistic tests we applied Kruskal-Wallis with Bonferroni post-hoc correction.

### 3.2 Relationship between understory plant species and canopy structural attributes

The DHP-based clusters were significantly different regarding species richness (Kruskal-Wallis test, χ2 = 13.92, p= 0.003) and Shannon diversity index (Kruskal-Wallis test, χ2 = 10.36, p= 0.01) of understory plants (Figure 5). The DCP-based clusters, however, differed significantly only regarding the species richness of understory plants (Kruskal-Wallis test, χ2 = 6.95, p= 0.03) (Figure 5). The larch and spruce-dominated clusters 4 and 3 of DHP showed to have the most plant species in the understory compared to the beech-dominated cluster 1 (Post-hoc test with Bonferroni correction, p<0.05). The same was true for the DCP clusters, the larch-dominated cluster 3 had a higher species richness than the beech-dominated cluster 1 (Post-hoc test with Bonferroni correction, p<0.05).

**Figure 5.**
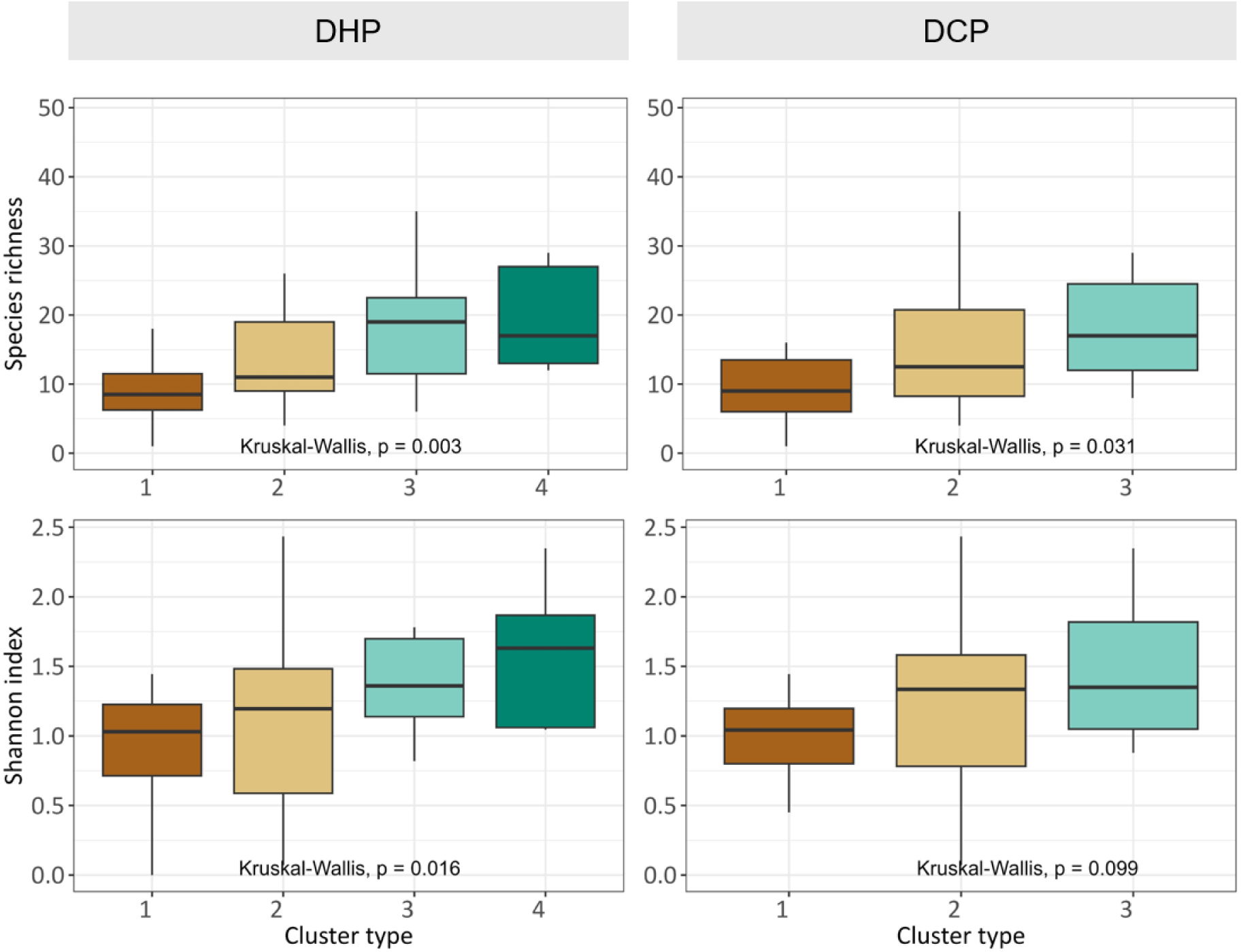
Kruskal-Wallis test and post-hoc tests with Bonferroni correction showing statistical differences between the four DHP cluster groups (left) and the three DCP cluster groups (right) regarding species richness (top) and Shannon diversity (bottom) of understory vascular plants.

Results from the canonical correlation analysis (Figure 6) shows what canopy structural attributes have the highest relative importance in explaining the understory plant composition. The most effective canopy structure attribute for explaining understory plant species variation in our forest sites was the clumping index derived from both photographic methods, DHP and DCP. The second most effective canopy structure attribute was mean leaf inclination from DHP. A Monte Carlo permutation test with 499 random permutations showed that all three parameters significantly explained the plant species variation, each explaining around 3.5% (pseudo-F = 1.6, 1.9, p-value = 0.002). The first two CCA axes had a cumulative explained variation of around 12%.

**Figure 6.**
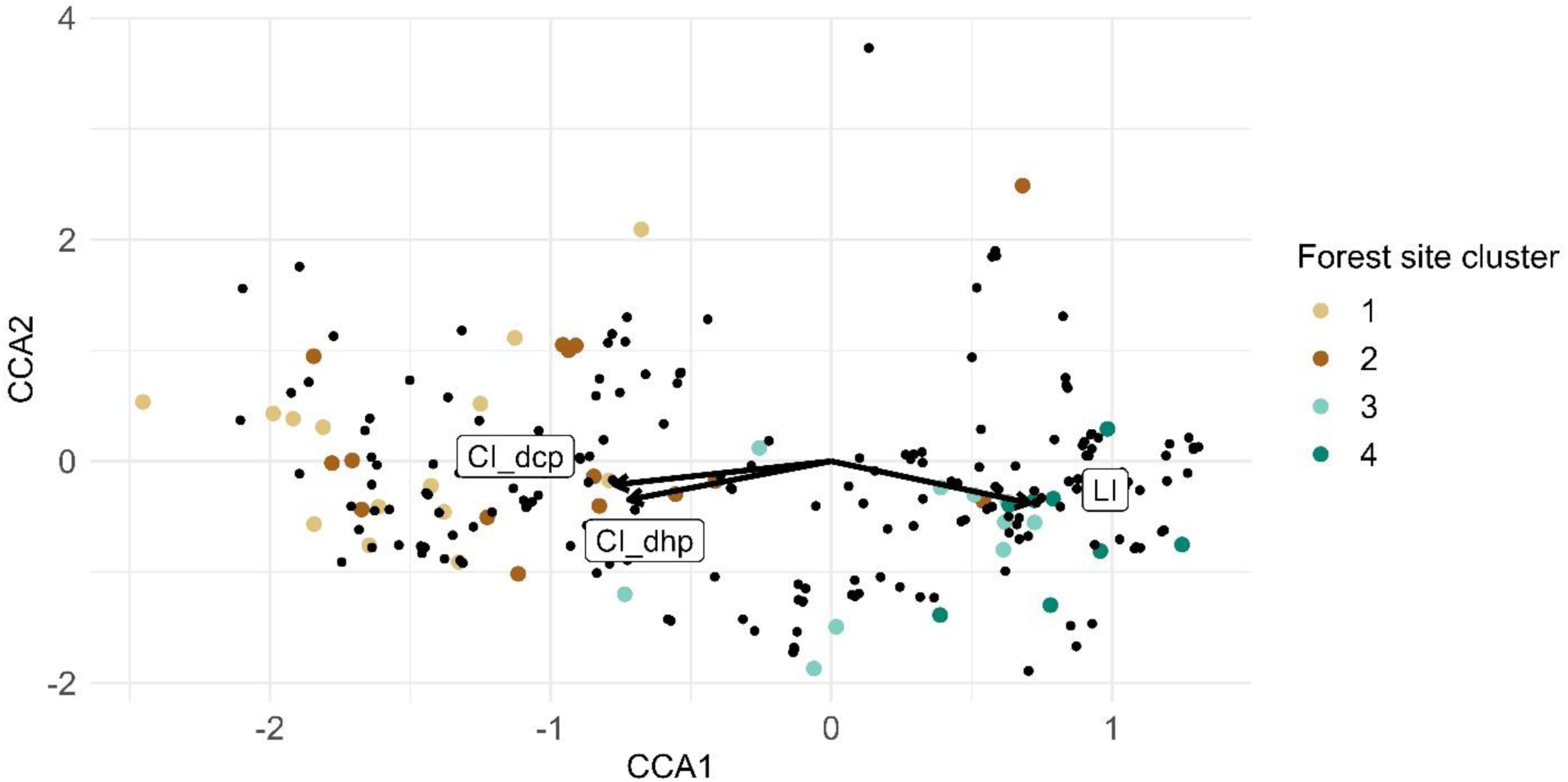
Canonical correlation analysis with the selected DHP and DCP canopy structure elements, clumping index (CI) and leaf inclination (LI), as vectors, and herbaceous species as black points. Forest sites were coloured according to DHP-based clustering (1 to 4; see Figure 3a). The length of the arrows explains the relative importance of the canopy structural elements. The forest sites and herbaceous species are placed together according to their relative similarity based on the selected structural elements.

DCP clumping index differed significantly between the four forest type clusters (Figure 7, Kruskal-Wallis test, χ2 = 27.9, p< 0.001). For example, the larch-dominated cluster 4 had the highest canopy clumping (i.e., more heterogeneous crown cover) while the beech-dominated cluster 1 the lowest clumping (i.e., more homogenous crown cover).

**Figure 7.**
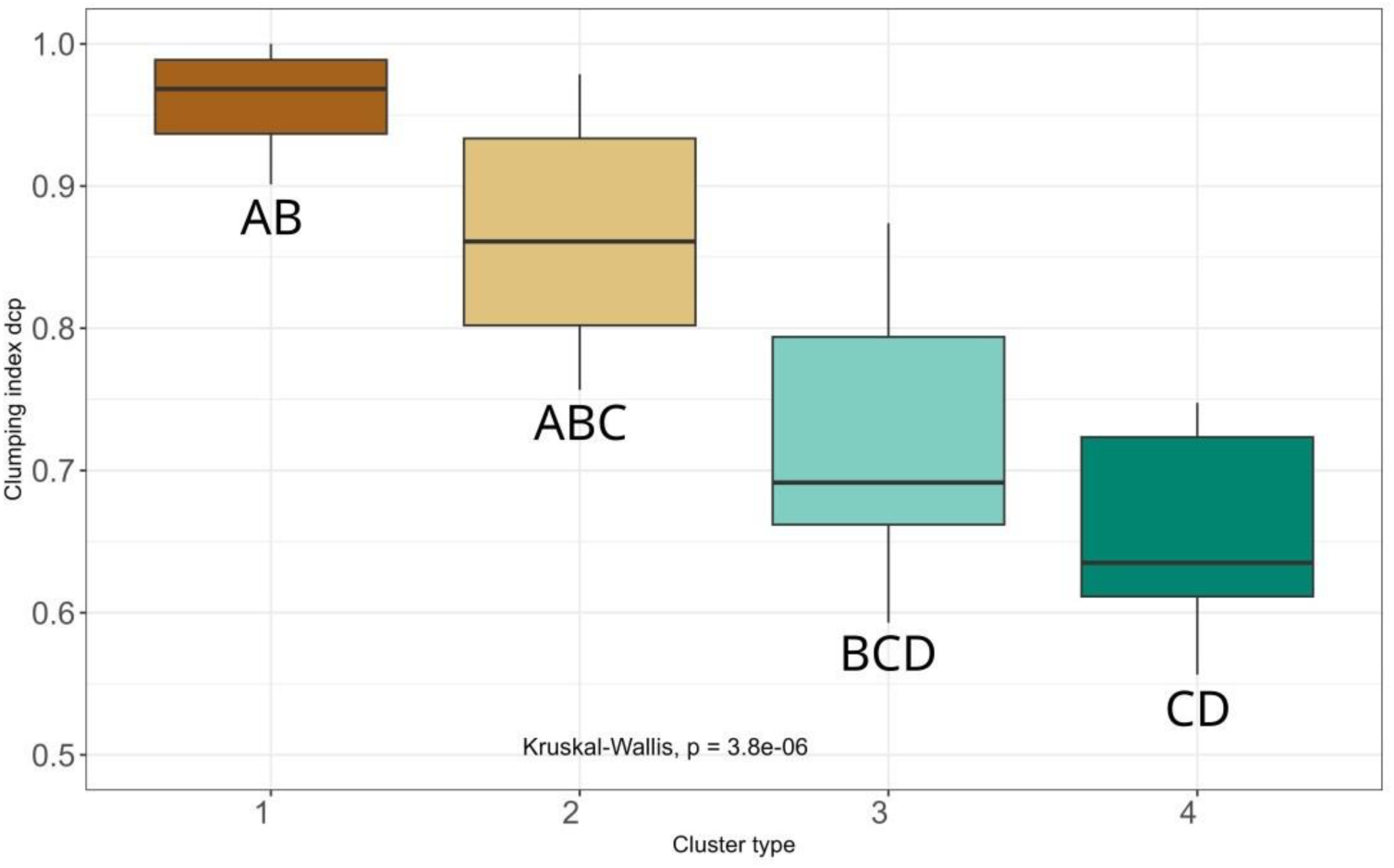
Boxplots to compare the DCP clumping index between the clusters based on the k-means clustering analysis done with DHP canopy structure. Letters above/below the boxplots indicate groupings according to post-hoc tests with Bonferroni correction.

## 4. Discussion

In line with our first hypothesis, our outcomes revealed that, between the two canopy photography techniques, hemispherical photography is most suitable to capture differences between forest types. The two most important canopy attributes to differentiate forest types were leaf inclination angle and canopy openness. We attributed the results as fisheye sensors integrate measurements of radiation transmittance at the full hemispherical view range (i.e., from different angles on the light transmission), while restricted view angle methods require independent measurements of leaf inclination angle, which cannot be estimated directly from DCP photos (Chianucci et al., 2018). Therefore, DHP allows to characterize a larger footprint of the canopy, being more representative of the light regime at the plot or stand level, while also directly determining the leaf inclination angle distribution. Previous studies have showed that leaf inclination angle is the result of different tree species ecological strategies, and there are compiled datasets showing how this attribute varies according to different species (e.g., Chianucci et al., 2018). Leaf inclination angle was shown to be an effective attribute to broadly differentiate between tree species, especially for broadleaf species (Falster and Westoby, 2003; Pisek et al., 2022). For conifer species, instead, there was a lack of empirical evidence that methods based solely on canopy photography can effectively be applied to differentiate conifer forests according to their dominant species. To date, methods for acquiring leaf inclination angle in conifers stands focused on photography obtained from above the canopy or laboratory research as opposed to field research with a fisheye lens (Yan et al., 2021). Our study shows that DHP-based canopy attributes can also distinguish among different conifer forests, as it was the case between our spruce/fir-dominated sites (DHP cluster n.3) and larch/Swiss stone pine sites (DHP cluster n.4).

Canopy openness derived from hemispherical sensors was also found to be a valuable attribute to differentiate forest types. It is known that light-demanding tree species typically form sparse canopies whereas the late successional or shade tolerant species have a denser canopy (Canham et al., 1999). This is a common ecological feature as light demanding species are usually present in the earlier stages of succession, setting the stage for late successional species to take over (Valladares and Niinemets, 2008). In our analysis, the variable canopy openness helped differentiate light-demanding subalpine forests (i.e., dominated by European larch and Swiss stone pine) from spruce-fir or beech dominated forests, which featured a denser canopy typical of late successional forest stands. Combined with other attributes such as leaf inclination angle, canopy openness can therefore be used as meaningful indicator to categorize forest research sites, which can be particularly useful in forest ecology studies featuring a multitude of stand types across a large environmental gradient (Rigo et al., 2024). Conversely, foliage and crown cover, which are the main attributes derived from DCP, are measures that reflect how dominated a forest site is by trees (Jennings et al., 1999; Ryu et al., 2012). This has less ecologically-relevant information – in this line, it is not surprising that only LAI values are the most effective attributes in DCP for discriminating forest types, as different forest types may display different LAI values depending on tree species features (Parker, 2020). In general, results from this study indicate that DHP has more potential in forest ecology and biodiversity studies, while DCP could be more suited for forest inventory (Salas-Aguilar et al., 2017), remote sensing (Cuba et al., 2018; Toda et al., 2022) or continuous tree monitoring and phenology (e.g. Chianucci et al. 2022).

When looking at the richness and alpha diversity (i.e., Shannon diversity index) of understory vascular plants, as expected, our findings showed that these two variables are naturally higher in more open forests (Van Couwenberghe et al., 2011). This can be observed in our sites too, such as those with a higher canopy openness in DHP-based cluster 4 and those with a lower LAI in DCP-based cluster 3. Both photographic methods used to cluster forest sampling plots were found to be appropriate to differentiate forests in terms of species richness of understory vascular plants, but only DHP-based clustering showed significant differences for alpha diversity. This can be explained because DHP is better suited to capture not only the quantity of light reaching the forest floor – expressed mainly by the LAI as the main attribute used to group forest sites with DCP – but also the quality of light reaching the understory – expressed by both canopy openness and leaf inclination in DHP-based clusters. This means that the diversity of understory plants is not only related to the amount of understory light penetrating through the canopy but also to the diversity of gap structures (i.e., crown heterogeneity, large vs small gaps) that create different microhabitat conditions linked to a higher diversity (Horváth et al., 2023).

In contrast to our second hypothesis, leaf clumping rather than canopy openness was the most important attribute for determining plant species distribution of the understory. Other research mentioned canopy openness as the most important factor (Ádám et al., 2018; Hederová et al., 2023; Lu et al., 2013; Yu and Sun, 2013). The fact that clumping index better explained the variation of the understory plants can be once more attributed to the fact that it is linked with the heterogeneity of light at the forest floor. The clumping index indicates in what way the leaves of the canopy are distributed (see graphical example and explanation in Figure A2). In general, sparser canopies exhibit more clumped distribution of foliage, which can be attributed to a larger frequency of large gaps (Chen and Cihlar, 1995) and a larger variation in gap size occurring at increasing canopy space availability (Nilson, 1999). Conversely, denser canopies showed a lower clumping, with more randomly distributed foliage (Liang et al., 2023), probably because the lower number of small gaps occurring at saturating canopy density (Macfarlane 2011). While both openness and clumping determine understory structure, the former is directly related to the total amount of light availability at the forest floor, while clumping also indicates how light is heterogeneously distributed. The importance of this canopy attribute could perhaps be credited to the fact that variation of light introduces different niches and thus attracts diverse herbaceous species. A modelling study by Kim et al. (2011) also showed that including the clumping of shoots in the model decreased the total amount of absorbed light by the canopy by 40%, thus with more clumping of the shoots more light falls to the forest floor. This is a substantial extra amount of light which could be an explanation why clumping resulted an important factor in our results. Hence, clumping is potentially more linked with diversity as diverse gap structures can create different microclimate conditions enhancing diverse plant species with different ecological strategies. This knowledge can have practical implementation in managed forests where specific interventions, such as selection harvesting promoting heterogeneous light conditions, can be applied to increase the diversity of understory community (Chianucci et al, 2024; Kovács et al., 2018), which in turn is an essential component of ecological resilience in forest landscapes (Messier et al., 2019).

While we here presented novel outcomes for the research field, we also recognize some limitations in our approach that future development could address. One shortcoming in our dataset was that in some research sites the botanical survey and the canopy photography were not gathered within the same year. However, as no recent silvicultural interventions or gap-opening natural disturbances were recorded during the monitoring period (2019-2023), it is unlikely that the changes in canopy structural attributes from one year to the next had a major influence in our results (i.e., a slight increased canopy closure due to growth). Another limitation was that the botanical survey of understory plants was executed in more detail starting from two subplots situated in the upper and lower corners of the research plot. We correlated this data with canopy attributes estimated as averages of the five – for DHP – or nine – for DCP – photo points distributed across the research plot. This could have influenced the significance of the relationship between overstory and understory, which might be one of the reasons why canopy attributes derived from DHP, capturing a largest footprint of the canopy, were found to be more suitable than DCP for relationship with understory plant diversity. This limitation could have been overcome with better harmonized sampling design of understory plants (i.e., by distributing botanical subplots in multiple sections within the entire plot), which however was unfeasible in the framework of the resources for this study. Another aspect that might have influenced our results was that the angle of view used to derive the canopy openness was 180 degrees, while Hederová et al. (2023) found that canopy openness derived from an 80 degrees angle of view yields stronger results in explaining the variability of understory vascular plant species. It is possible that the angle of view used in our study was not appropriate enough to fully capture the canopy structure. The fact that the percentage explained variance was fairly low could be expected from other literature indicating that other key factors influencing the abundance and diversity of understory plants, such as understory temperature, soil pH, and humidity (Díaz-Calafat et al., 2023; Ewald, 2000; Schauer et al., 2023). We acknowledge that including such factors would have likely increased the cumulative explained variation of understory plant diversity, but this would have deviated from the aim of the study focusing on canopy structural attributes from different photographic techniques. Purposely omitting other environmental factors also allowed us to demonstrate leaf inclination as a driving factor of the variation in the understory plant species, as it was also a distinguishing feature of forest type. This suggests that this canopy attribute could serve as an indicator for swiftly differentiating between forest types without necessarily using on other in-situ data that require significantly more effort to collect (i.e., full tree callipering).

## 5. Conclusions

Our study demonstrates that canopy structural attributes derived from digital photography – in particular hemispherical photography – can be used as meaningful indicators for forest biodiversity research and monitoring. Using canopy attributes such as leaf inclination and canopy openness – which can be derived relatively rapidly in the field compared to more traditional field sampling methods – could be used as meaningful indicator to categorize forest research sites in forest ecology and biodiversity research, particularly for investigations across a multitude of stand types in terms of structure and composition. Canopy photography methods could be also applied in forest management and planning, for example for rapid but precise estimates of forest cover at stand level which are often required for planning small-scale silvicultural interventions.

This study also shows that the understory plant communities were more impacted by the heterogeneity of horizontal canopy structure, expressed by foliage clumping, rather than other attributes related simply to the amount of light or foliage such as openness or LAI. This is interesting and useful knowledge in the context of management regimes in forests of high conservation value. For example, silvicultural interventions based on single-tree selection could aim at increasing canopy heterogeneity, promoting a more diversified light regime at the forest floor and creating a light environment with a richer and more diverse understory plant community that can be beneficial for other forest dwelling taxa (e.g., insects, birds, mammals). If strategically planned across forest landscapes, such interventions could be beneficial for overall ecosystem functioning and ecological resilience.

## Author contributions

**Mina Marco**: Conceptualization, Funding acquisition, Project administration, Investigation, Methodology, Supervision, Writing – review & editing. **Chianucci Francesco**: Conceptualization, Investigation, Methodology, Writing – review & editing. **Anouk von Meijenfeldt**: Conceptualization, Data curation, Methodology, Investigation, Formal analysis, Visualization, Writing - original draft, Writing – review & editing. **Francesca Rigo**: Investigation, Writing – review & editing. **Jente Ottenburghs**: Supervision, Methodology, Writing – review & editing. **Andreas Hilpold**: Resources, Project administration, Funding acquisition, Writing – review & editing.

## Acknowledgements

We thank Abraham Mejia Aguilar for kindly allowing the use of his photographic equipment, Simon Stifter and Lisa Angelini who collected the understory plant data, Sebastian Marzini for the support in the field and visualization, and Luca Frattini for the help in preparing the botanical data.

## Data Availability

If required, the data sets supporting the results of this article (values of canopy structure attribute for each plot) will be made available in the Zenodo repository upon acceptance.

## Appendix

**Figure A1.**
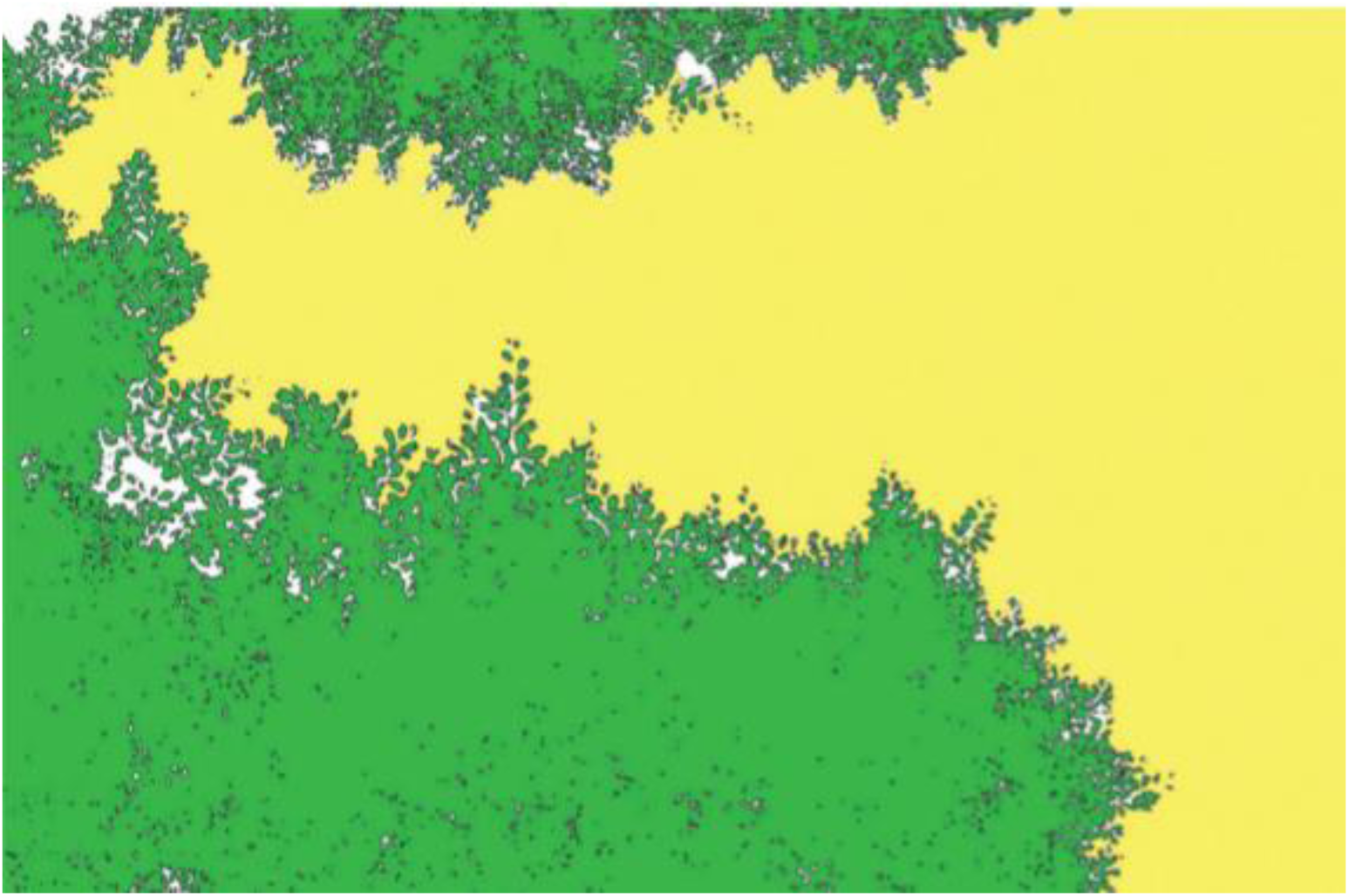
An example of a DCP image, which has been classified in large between-crowns canopy gaps (yellow), and small, within-crown canopy gaps (white). Foliage cover (green) is the fraction of pixels that does not lie in canopy gaps, while crown cover is the fraction of pixels that does not lie in between-crowns gaps.

**Figure A2.**
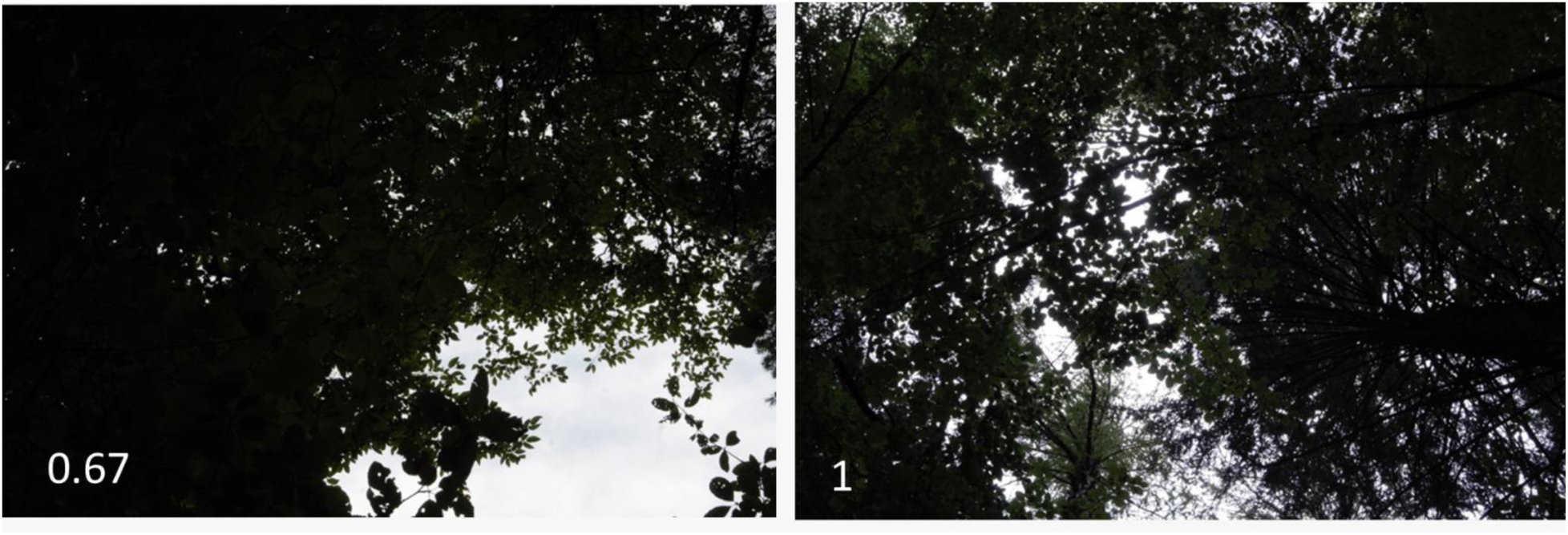
Example of canopy clumping. Two DCP photographs in the same forest site but on two different photo points. The left one with a high clumping (i.e., more aggregated distribution of foliage and more large openings; values of CI between 0 and 1). The right one has a low clumping (i.e., more random distribution of foliage and more homogeneous openings; values of CI close to 1).

**Table A1.**
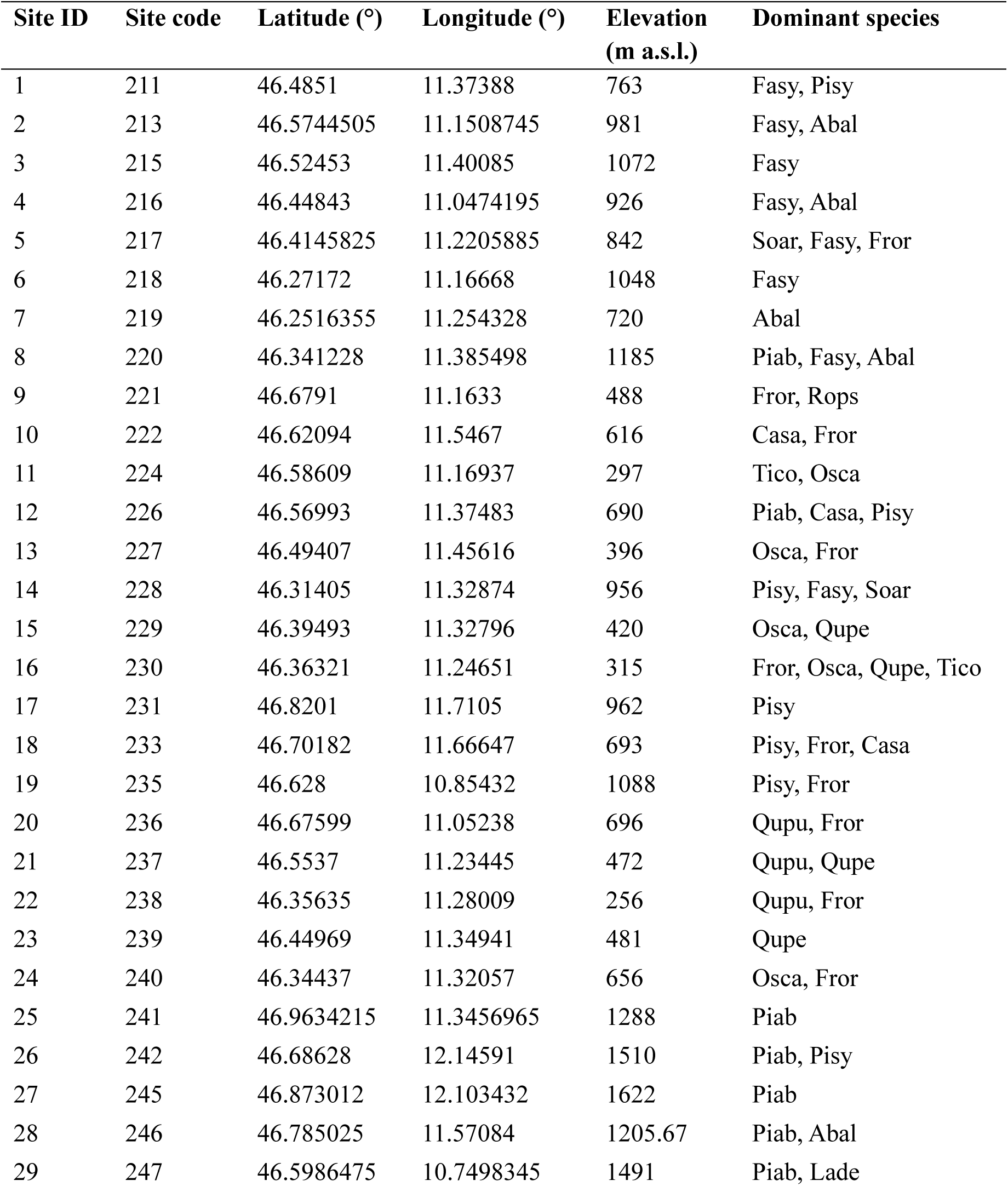
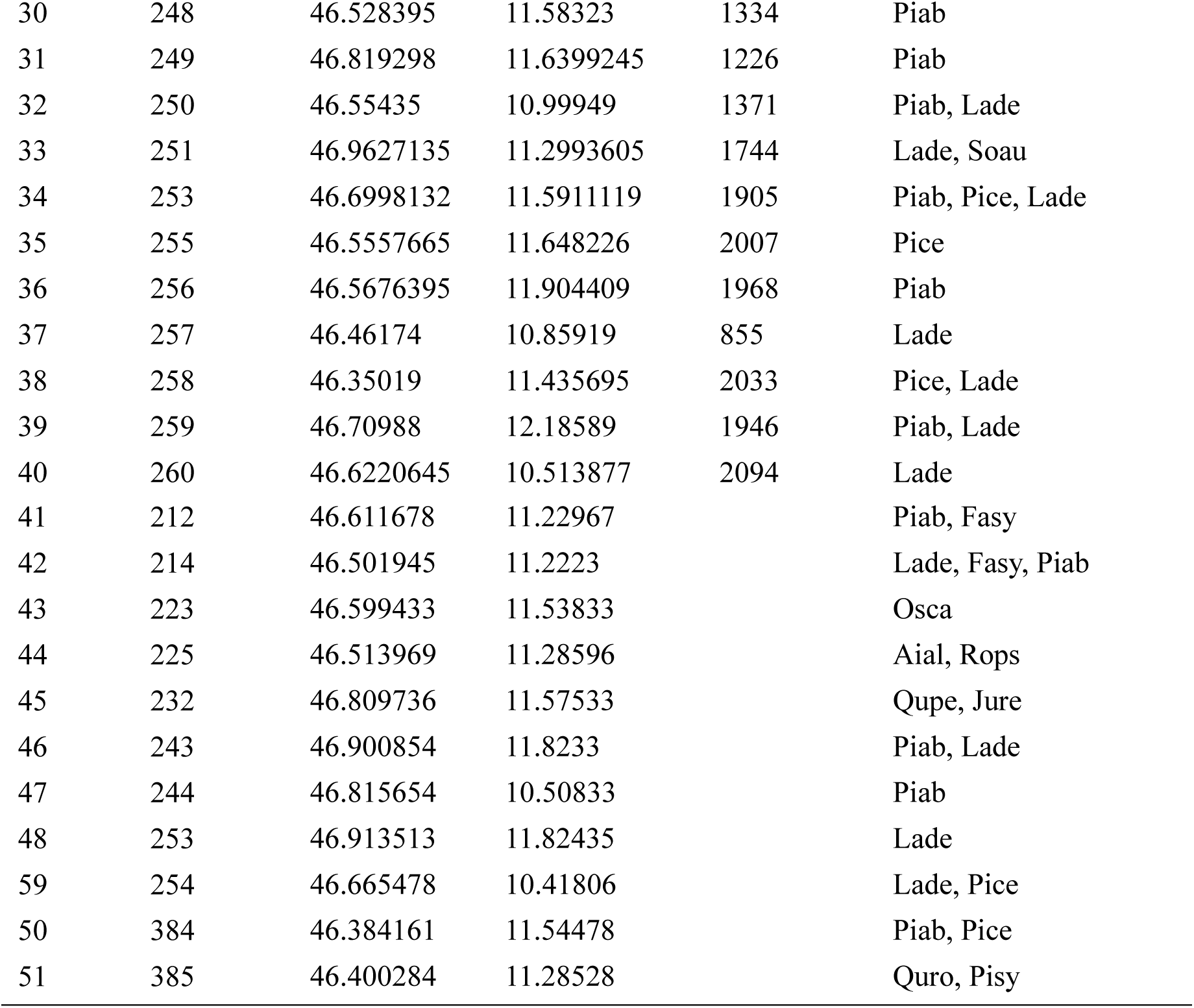
Location of the 51 forest sites with their serial ID and site code used in the biodiversity monitoring project, along with their elevation and the list of dominant tree species (Fasy = *Fagus sylvatica*, Abal = *Abies alba*; Piab = *Picea abies*; Fror = *Fraxinus ornus*; Casa = *Castanea sativa;* Tico = *Tilia cordata*; Osca = *Ostrya carpinifolia*; Pisy = *Pinus sylvestris*; Qupu = *Quercus pubescens*; Qupe = *Quercus petraea*; Lade = *Larix decidua*; Soar = *Sorbus aria*, Pice = *Pinus cembra*; Soau = *Sorbus aucuparia*; Rops = *Robinia pseudoacacia*; Aial = *Ailanthus altissima*; Jure = *Juglans regia*).

**Figure A3.**
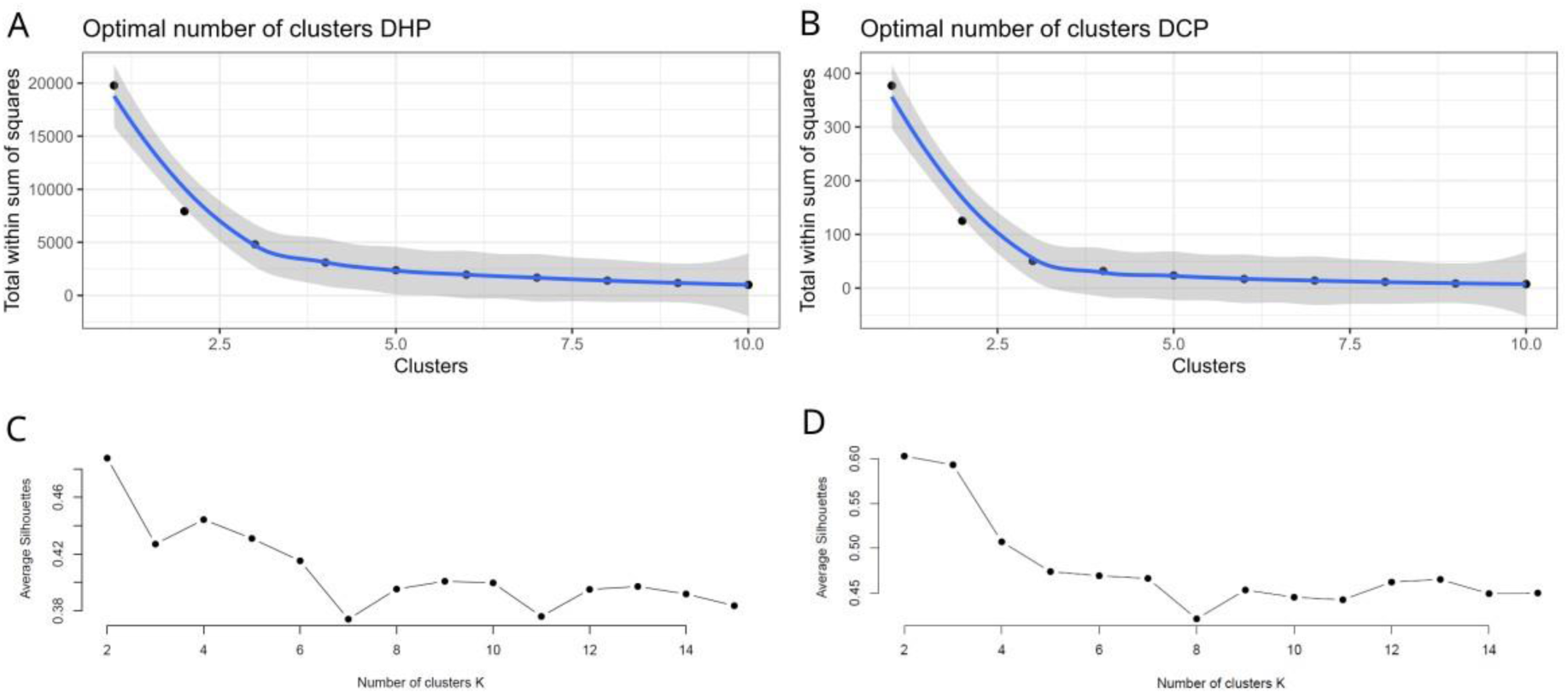
Total sum of squares and silhouette test of cluster analyses to find optimal number of clusters in the DHP (panel a and c) and DCP (panel b and d) PCA’s.

**Figure A4.**
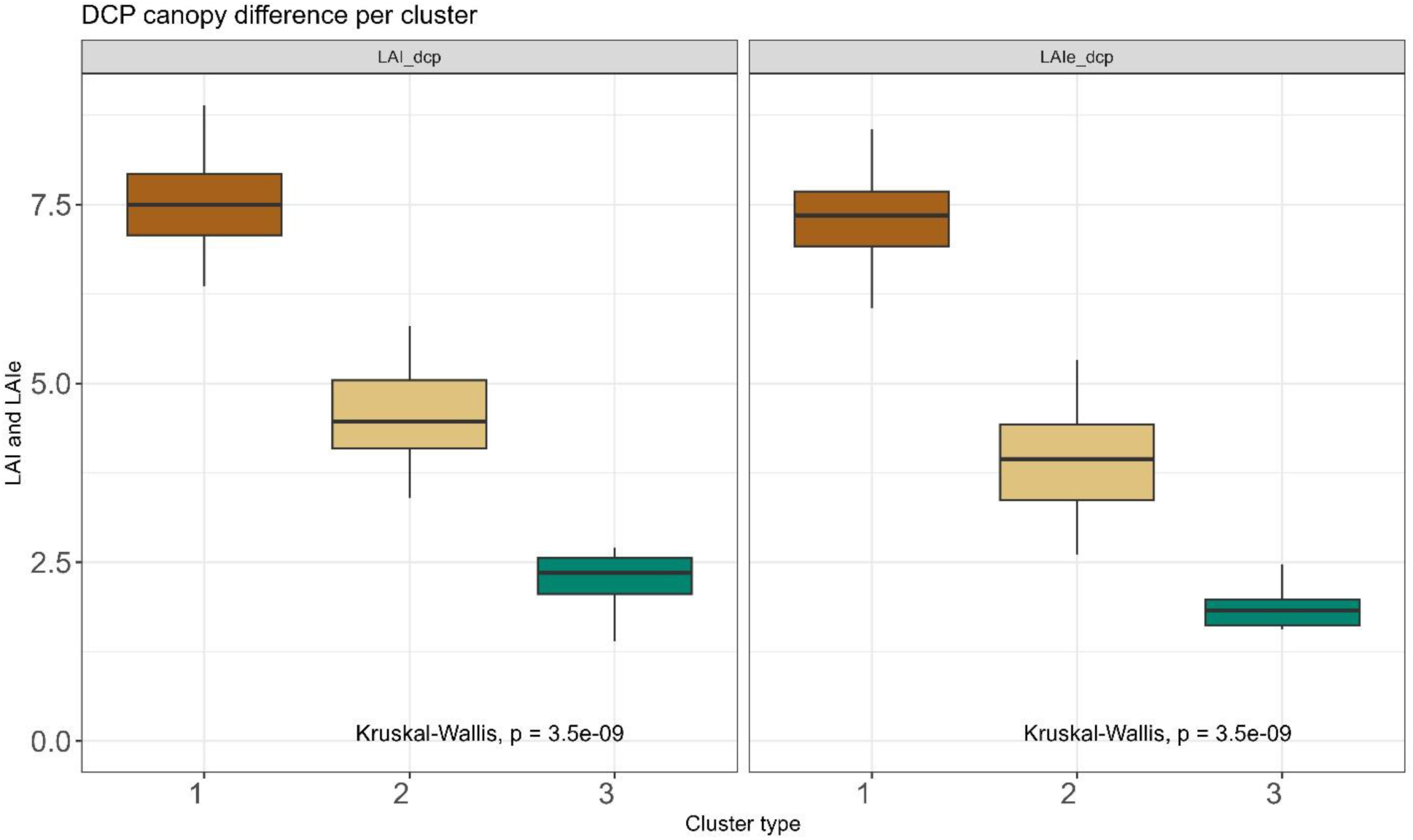
Kruskal-Wallis test and Bonferroni post-hoc showing statistical differences between the three DCP cluster groups for the two canopy attributes actual LAI (left) and effective LAI (right).

